# Units of motor production: Bengalese finches interrupt song within syllables

**DOI:** 10.1101/2020.02.19.956698

**Authors:** D. Riedner, I. Adam

## Abstract

Birdsong consists of syllables that are separated by silent intervals. Previous work in zebra finches showed that syllables correspond to the smallest motor production units (Cynx, 1990; Franz and Goller, 2002) by inducing song stops using strobe light. In this study, we interrupted the song of six Bengalese finches experimentally with the bird’s own song as auditory stimulus using an interactive playback approach. Five of the tested males interrupted their ongoing vocalizations within syllables (16 instances of induced interrupted syllables) in response to the playback. Additionally, we observed 9 spontaneous interruptions in our control recordings. This study establishes that birds can interrupt ongoing syllables within extremely short latencies in response to an auditory stimulus, and that auditory stimuli interrupt syllables more effectively than visual stimuli. Even if syllables are the functionally stable production units, the ability to disrupt those units differs between species and individuals, indicating various degrees of vocal control.

## Introduction

Birdsong is composed of series of syllables which occur in a regular pattern forming a motif and are separated by silent intervals. To determine the motor production unit of birdsong, previous studies (Cynx, 1990; Franz and Goller, 2002) used strobe light as a disruptive stimulus. In the vast majority of cases, interruptions occurred during silent intervals between syllables (Cynx, 1990) at the end of expiratory pressure pulses (Franz and Goller, 2002). These findings indicate that syllables, are the smallest motor production units (Franz and Goller, 2002). An additional study (Seki et al., 2008) also used strobe light to show that stereotypic syllable sequences in Bengalese finches (*Lonchura striata var. domestica*) were more order dependent and harder to interrupt than variable sequences, suggesting the existence of a second-order unit of motor control organization. Studies with the same methods and similar outcome were conducted in nightingales (*Luscinia megarhynchos*) (Riebel and Todt, 1997), in non-songbirds, collared doves (*Streptopelia decaoto*) (tenCate and Ballintijn, 1996), and in cotton-top tamarins (*Saguinus oedipus*) (Miller et al., 2003). These studies indicate that individuals can exert control over syllables in a vocalization, suggesting that the basic motor production unit of the song presumably correspond to syllables.

## Objective

We aimed to investigate the impact of an auditory stimulus on ongoing vocalizations. We analyzed whether experimental interference using the bird’s own song (BOS), led to singing interruptions. Our expectation was that ongoing production of a syllable cannot be interrupted, given that syllables are thought to be the smallest motor production unit.

## Results and Discussion

Interruptions within syllables were observed in five out of six Bengalese finches (Figure 1). In total, we successfully targeted 136 syllables with playbacks (range: 21-25, average: 22.6 ±1.5 per bird) of which 11.7% (16 syllables) led to an interruption of the targeted syllable (range: 0-34.7 %, average: 11.5 ± 12.6 per bird) (Figure 1B, D). Additionally, we observed 9 spontaneous interruptions in 2 males during the control period (50 minutes of song in for 6 birds, per bird: mean 493±96s, range 370-638s) (Figure 1E). Interruptions did not seem to occur at a specific position within a bird’s motif. The likelihood to interrupt an ongoing syllable differed between animals. The two males, which had the highest percentage of interrupted syllables in the playback experiments (Ls1 and Ls2) were the only two males that spontaneously interrupted syllables in the control period.

**Figure 1:**
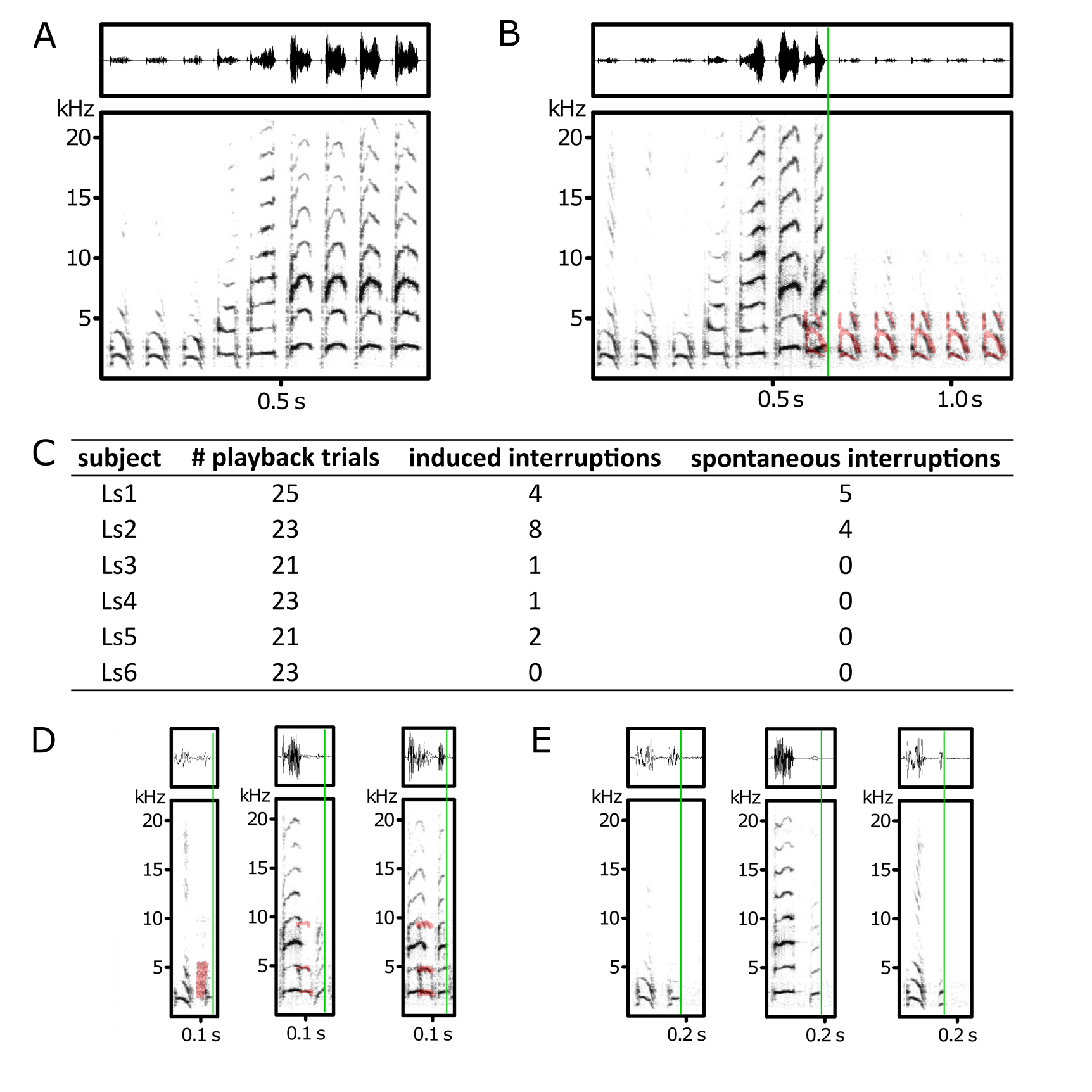
Bengalese finches can interrupt ongoing syllable production in response to auditory input as well as spontaneously. A) Example of uninterrupted song of a Bengalese finch. B) Example of an induced syllable interruption (indicated by the green line) caused by targeted BOS playback (highlighted in red). C) Summary of the number of playback trials, induced and spontaneous syllable interruptions for each subject. Examples of induced (D) and spontaneous (E) syllable interruptions.

In previous studies, using strobe light to interfere with singing activity (Cynx, 1990; Riebel and Todt, 1997; Franz and Goller, 2002; Seki et al., 2008) interruptions occurred after completion of the ongoing vocalization. Furthermore, electrophysiological stimulation of the robust nucleus of the archistriatum (RA) and HVC (used as a proper name), both part of the song system, leads to perturbations of song (Vu et al., 1994), indicating that syllables rather than the whole song correspond to motor production units. However, within anecdotal syllable interruptions have been reported in all published studies. Strobe light induced three within syllable interruptions in two zebra finches (Cynx, 1990) and (Seki et al., 2008) also reported that song stops sometimes occurred during the ongoing syllable in Bengalese finches when interrupted with strobe light. However, to our knowledge a systematic description and quantification of interrupted syllables has not been conducted so far.

In our experiments we used an auditory stimulus (BOS) to interfere with sound production, instead of strobe light. We assumed that auditory, naturally occurring stimuli would be more relevant for the behavioral repertoire of songbirds and less stressful than artificial ones like strobe light or white noise. Compared to the previous studies (Cynx, 1990; Riebel and Todt, 1997; Franz and Goller, 2002; Seki et al., 2008), induced syllable interruptions occurred in the majority of tested subjects. The wide range of susceptibility to induce interruptions between males suggested that the degrees of vocal control differed between subjects. Even though quantitative data is mostly lacking, syllable interruptions in response to auditory stimuli in our experiments seemed to be more frequent than in zebra finches induced by strobe light (Cynx, 1990), which suggested that behaviorally more relevant auditory stimuli are more effective than a visual stimuli. This finding is supported by a study in cotton top tamarins in which auditory stimuli (white noise) were also more successful in interrupting vocalizations than visual stimuli (Miller et al., 2003).

Using our playback paradigm to induce singing interruption in zebra finches yielded no syllable interruption in 5 subjects. These results in combination with previously published data indicate that different songbird species also differ in their degree of control over ongoing vocalizations. Bengalese finches seem to have a greater degree of vocal control than zebra finches as almost all of our subjects interrupted ongoing syllables, while none of our subjects and only two of seven tested zebra finch males interrupted syllables in a previous study (Cynx, 1990).

In conclusion, Bengalese finches can interrupt ongoing syllable in response to sensory stimuli, which raises the question whether syllables really are the functionally stable production unit. Additionally, vocal control in response to sensory stimuli differs between individuals and species.

## Methods

### Animals song recordings

Six male Bengalese finches (*Lonchura striata var. domestica*) and five zebra finches (*Taeniopygia guttata*) >100 days post hatch, were used for song recordings and playback experiments. All subjects were kept in mixed sex group aviaries on a 13:11 light/dark schedule and provided with food and water ad libitum. All experiments were conducted in accordance with the Danish law concerning animal experiments and protocols were approved by the Danish Animal Experiments Inspectorate (Copenhagen, Denmark).

### Song recordings and selection of control songs

For song recordings and playback experiments birds were kept in custom built sound attenuated chambers (60 x 95 x 57 cm). Each chamber was equipped with an omnidirectional microphone (Behringer ECM8000) positioned centrally 12 cm above the cage. Vocalization activity was monitored and recorded continuously with sampling frequency of 44.1 kHz by a standard desktop computer running Sound Analysis Pro (Tchernichovski et al., 2000). Recordings of undirected song (Sossinka and Boehner, 1980) were started 24 hours after a bird was transferred to the recording chamber to allow for habituation to the new environment. Each bird was recorded for three days as control period. The first 30 song bouts within the three days were used as control vocalizations. A bout was defined as singing activity with silent intervals smaller than 1.5s.

### Playback selection

For each animal 10 different playbacks were generated from each bird’s own song by selecting short sections (650 ms) of song from the control recordings using Avisoft-SASLab Pro 4.36 software (Avisoft Bioacoustics, R. Specht, Berlin, Germany). Playback files were high-pass filtered at 1 kHz. To ensure that the overall power of the playback files was equal, the root mean square (RMS) amplitude was normalized to a common value, using the seewave package (Sueur et al., 2008) in R version 3.4.0 (R Core Team, 2017)(R Core Team (2017), http://www.Rproject.org/).

### Playback experiment

We used a system to automatically deliver targeted playbacks of BOS based on the Teensy-TAF system by Hamish Mehaffey (http://github.com/whmehaffey/Teensy3.6TAF). Our version (Teensy-TAP) consisted of a Teensy USB Board, (PJRC, Version 3.6) connected to a custom-made circuit board. The system was connected to portable loudspeakers (Hama, E80), which were placed in front of the cage within the sound chamber. The Teensy-TAP setup enabled us to automatically target specific syllables with playbacks with very short delays. Each time a target syllable was detected, playback of one of the 10 playback files was trigged and subsequent trigger events were discarded for at least one minute. Playbacks were selected in random order and played back at 12 bits. For each bird the detection code was adjusted to only trigger once per song bout and prevent triggers in silent periods. We considered each presentation of one playback as one trial and one session consisted of eleven to thirteen trials. Each male was subjected to two sessions, which were seperated by at least five days to reduce habituation. Before the second session started, the birds had one day to adjust to the sound chamber again. Although the system ran automatically, the experiments were monitored by an experimenter to ensure optimal triggering and to adjust the code if necessary.

### Data analysis

All sound files were high-pass filtered at 1 kHz to remove low-frequency background noise and converted into spectrograms (Avisoft-SASLab Pro, Hamming window, FFT-length: 512 samples, frame: 100%, overlap: 75%). Spectrograms of the control periods as well as the sessions were screened visually to identify and annotate song bouts, the playback (in session files) as well as interrupted syllables. We classified interrupted syllables in response to playback and in the control files as “induced” and “spontaneous”, respectively. The results were visualized using the ggplot2 package (Wickham, 2016)(Wickham, 2009) in R version 3.4.0 (R Core Team, 2017)(R Core Team (2017), http://www.Rproject.org/).

